# Perinatal high-fat diet induces systemic inflammation via altered tight junction complex organization during development and decreases expression of intestinal satiety peptides in male offspring

**DOI:** 10.64898/2026.11.12.766002

**Authors:** Lindsey K. Macias, Akul Kesarwani, Jason Perez, Jasmine Shakir, S. Andrew Aston, Zachary A. Cordner, Yuma Kitase, Lauren L. Jantzie, Claire B. de La Serre, Ken Y. Hui, Timothy H. Moran, Kellie L. Tamashiro

## Abstract

**Objective:** Epidemiological evidence suggests that maternal obesity increases risk for offspring obesity and comorbidities, pointing to the significance of critical periods of development. Adverse metabolic effects of high fat (HF) diet consumption on the gut microbiome and inflammation have been well documented in obesity models. Emerging evidence demonstrates that the gut microbiome may play a role in transgenerational obesity, however, underlying mechanisms still need to be elucidated to identify intervention targets.

**Methods:** A rat model was used to demonstrate whether maternal HF diet during gestation and lactation results in gut-brain axis deficits in rat offspring. Gene expression of small intestinal satiety hormones and C-fiber innervation in the nucleus of the solitary tract (NTS) were measured in adult offspring. At weaning, circulating inflammatory mediators and colonic microbial metabolites were measured in offspring.

**Results:** In adulthood, HF males had reduced vagal innervation in the NTS and reduced small intestinal expression of CCK, GLP-1, and PYY. At weaning, HF offspring had elevated levels of CRP and IL-1β and depleted levels of microbially produced colonic short chain fatty acids (SCFAs), which are critical for maintaining intestinal barrier function. HF males had reduced colocalization between tight junction proteins, Claudin-1 and ZO-1 at weaning, while HF females did not.

**Conclusion:** Disorganization of tight junction complexes in HF male offspring during development may play a role in the gut-brain axis deficits seen in HF males in adulthood. Together, these data demonstrate that inflammation-induced by HF diet can negatively impact gut-brain axis development and function.

## 1. Introduction

The prevalence of maternal obesity in the U.S. continues to rise with prevalence of maternal obesity and overweight in the U.S. currently at ∼60% [1]. Maternal obesity poses many health risks to the developing fetus and predisposes offspring to obesity, comorbid metabolic diseases, and mental health conditions [2–5]. The intrauterine and early postnatal environments are critical for proper development and growth of offspring. Preclinical data in animal models indicate that maternal high fat diet (mHFD) before and during pregnancy has a negative impact on fetal development and can predispose offspring to obesity and related metabolic diseases throughout life [6–10]. Understanding the physiological mechanisms induced by mHFD during development that may drive caloric overconsumption in offspring provides opportunity to develop novel strategies and early-life interventions to prevent obesity. We have previously demonstrated that mHFD in the perinatal period biases male but not female offspring toward obesity [10]. In this study, we test the overarching hypothesis that exposure to mHFD during the perinatal period increases offspring propensity for obesity by altering gut-brain axis structure.

### 1.1. Gut-brain axis controls of feeding behavior

Upon ingestion of food, the gastrointestinal (GI) tract is the first post-ingestional site of interaction between the host and meal, eventually informing the central nervous system about the size and composition of the meal. Throughout the GI tract are specialized enteroendocrine cells (EECs), that secrete hormones and peptides involved in satiety signaling to the brain in response to intestinal stimuli [11]. The vagus nerve serves as the primary neural connection that communicates meal related information from the gut to the brain. Vagal afferent neurons (VAN) respond rapidly to meal-related gut stimuli including sensory information like mechanical distension [12] macronutrient composition [13], and satiety peptide hormones including CCK [14, 15], PYY [16] and GLP-1 [16]. Post-ingestive feedback signals are relayed from the gut to the brain by sensory vagal afferent nerve fibers that ultimately terminate in the nucleus tractus solitarius (NTS) located within the brainstem. HF diet in adult rodents alters both the function and connectivity/structure of VAN, reducing vagal c-fibers in the NTS [17], impairing gut-brain communication and overall food intake control [18, 19].

### 1.2. Microbiota impact on gut-brain axis function

Emerging evidence has begun to identify the gut microbiome as a key player in modulating gut-brain axis controls of feeding behavior. Obesity is often associated with greater intestinal permeability (“leaky gut”), microbiota dysbiosis and low-grade inflammation [20]. HF feeding triggers rapid changes in gut microbiota composition or dysbiosis in humans and rodents [8, 17, 21–23]. Alterations include reduced amounts of bacteria that produce short chain fatty acids (SCFAs) and increased population of pro-inflammatory microbes [20, 21, 24, 25]. Overgrowth of pro-inflammatory-type microbiota is sufficient to cause weight gain [26–28]. Microbiota dysbiosis induced by HF diet can remodel vagal structure and reduce gut-brain axis communication of satiety signaling [18]. Adult rats fed HF diet have reduced vagal projections to the NTS and preventing dysbiosis prevents loss of NTS vagal innervation, preserving the satiety response to exogenous CCK [17, 18]. Colonization of adult germ-free (GF) rats with a HF-type microbiota, is sufficient to promote withdrawal of NTS c-fibers, overeating and weight gain [29]. While is it clear that HF-induced dysbiosis can alter gut brain axis controls of feeding behavior, what is less clear is the role of maternal HF diet on the developmental programming of gut-brain axis controls of feeding behavior.

Our previous data indicate that perinatal HF diet alters maternal care, milk composition and consumption, and offspring gut microbiota composition during the early postnatal period [8, 30, 31]. In this study, we first aim to address whether mHFD results in altered gut-brain axis development and function. We hypothesized that HF offspring have reduced vagal afferent innervation in the NTS and reduced expression of intestinal satiety peptides involved in meal termination. Secondly, we aim to identify potential mechanisms by which mHFD-induced gut-brain axis deficits occur. Based on our previous studies in which we demonstrate that HF offspring have a dysbiotic gut microbiome during a critical period of development [31], we predict that HF offspring also have decreased colonic SCFAs, disorganization of tight junction proteins, and increased systemic inflammation, which may be driving gut-brain axis deficits.

## 2. Materials and Methods

### 2.1. Animals

Cohorts of timed-pregnant Sprague Dawley rats procured from Charles River (Raleigh, NC) were received on gestation day (G)2. Animals were individually housed in tub cages in a temperature- and humidity-controlled room maintained on a 12-hour light/dark cycle with lights on at 0600. Dams were divided into 2 weight-matched groups fed either standard laboratory chow (CH) diet (Teklad 2018; 3.1 kcal/g, 18% kcal from fat, 24% kcal from protein, 58% kcal from carbohydrates) or a purified HF diet (Research Diets D12492, 5.24 kcal/g, 60% kcal from fat, 20% kcal from protein, 20% kcal from carbohydrates) during gestation and lactation (n=6-8/diet). Dams were weighed and food intake measured daily throughout gestation and lactation. The day of parturition (G22 or G23) was defined as postnatal day zero (P0). On P1, litters were culled to 10 pups per litter (5 male, 5 female) to control for litter size and pups’ access to nutrition. On P21, all pups were removed from dams and weaned to CH for the remainder of the experiment. Offspring were group housed until adolescence, at which point they were housed individually. The data here were collected over 3 cohorts of animals, each cohort with a sample size of n=6-8 litters per each diet group. Only one male and one female offspring from each litter was used for each individual experiment at each timepoint. All animal procedures were approved by the Animal Care and Use Committee of the Johns Hopkins University School of Medicine.

### 2.2. Tissue and Plasma Collection

Plasma and tissue samples were collected from representative pups from each litter (1 pup/sex/litter) at two timepoints, weaning at P21 and in adulthood at P120. Blood was collected from male and female pups by rapid decapitation in EDTA tubes and centrifuged at 3,000g for 15 min at 4°C. Plasma was stored at -80°C and then used to measure inflammatory markers. For the small intestinal gene expression measured at P21 and P200, 2cm segments of tissue were collected from the duodenum, jejunum, and ileum, flushed with PBS with 0.02% sodium azide, and immediately snap frozen and stored at -80°C. Distal colons were also collected at P21 for quantification of tight junction proteins using western blot or immunofluorescence. Colonic tissues used for western blot were flushed with PBS with 0.02% sodium azide and immediately snap frozen for storage at -80°C, while colonic tissue used for immunofluorescence and histology were flushed with PBS, and immersion fixed in 10% neutral buffered formalin for 24-hrs prior to routine tissue processing and paraffin embedding.

### 2.3. Plasma Leptin

Plasma samples were isolated from blood collected from pups at weaning on P21 immediately following rapid decapitation, EDTA treatment, and centrifugation. Samples were analyzed using an enzyme-linked immunosorbent assay (ELISA) for rat leptin (Sigma-Aldrich EZRL-83K) in accordance with manufacturer instructions. ELISA plates were read on SpectraMax i3x using SoftMax Pro GXP microplate analysis software (Molecular Devices). All samples were run in duplicate with average intra- and inter-assay coefficients of variations (CVs) ≤ 10%.

### 2.4. NTS Immunohistochemistry

Immediately following euthanasia, rat offspring (1 pup/sex/litter) were transcardially perfused with 0.1 M phosphate-buffered saline (PBS; pH 7.4) followed by 4% paraformaldehyde. Hindbrains were dissected and submerged in 4% paraformaldehyde for 24 hours, followed by 30% sucrose in PBS and 0.1% NaN3 (Sigma-Aldrich; pH 7.4). Hindbrains were frozen to -20°C and cryosectioned at 20 µM thickness (Leica CM3050S) from the caudal to rostral direction to capture the NTS between bregma coordinates -14.16 to -12.93 mm as previously described [18] . Sections were slide mounted and stained for isolectin B4 (IB4) using a biotin conjugated IB4 (Vector Laboratories B-1205) to visualize unmyelinated c-fibers. Slides were first incubated in 0.1% Triton X in PBS at room temperature for 10 min, washed with phosphate buffered saline (PBS) for 10 min, and blocked with 20% goat serum in PBS for 1 hr at 37°C in a humidified chamber. Slides were incubated overnight at 4°C in a humidified chamber with 1:100 IB4 and 1% goat serum in PBS. A negative control slide with no added primary antibody was also included to account for nonspecific background staining. All slides were thoroughly washed in PBS for 10 mins and incubated in a secondary antibody solution with 1:500 ExtrAvidin-Cy3 (Sigma E4142) and 1% goat serum in PBS in a humidified chamber at 37°C for 1 hr. Slides were again thoroughly washed with PBS for 10 mins prior to coverslipping with Vectashield mounting medium with DAPI (Vector Laboratories H-1200-10). Slides were sealed with clear nail polish and stored at 4C. Images were captured using Zeiss AXIO Imager.D1 at 4x magnification. IB4 fluorescence positive staining was quantified at the level of the area postrema using ImageJ software as previously described [18].

### 2.5. Gene Expression of Intestinal Satiety Peptides

Cholecystokinin (*Cck*), glucagon like peptide 1 (*Gcg*), and peptide YY (*Pyy*) expression levels were assessed relative to beta-actin (*Actb*) throughout the small intestine using quantitative real-time PCR (RT-qPCR) in adult offspring as measures of enteroendocrine cell (EEC) function. Total RNA was extracted using the RNeasy Mini Kit (Qiagen). cDNA was generated using the QuantiTect Reverse Transcription Kit (Qiagen). RT-qPCR reactions were carried out in triplicate using 1xTaqMan master mix (Applied Biosystems), 1xTaqMan probes for each gene (Supplementary Table 1), and 2 μg of cDNA in a total of 20 μL. Real-time PCR reactions were performed on an Applied Biosystems 7900HT Fast Real-Time PCR system under standard conditions for 40 cycles. Gene expression data were analyzed using the –ΔΔCT method. Gene expression levels were then normalized relative to the CHOW duodenum, the average of which was set equal to 1.

### 2.6. Plasma C-Reactive Protein (CRP)

Plasma C-reactive protein (CRP), was measured in plasma collected at P21 by enzyme-linked immunosorbent assay ELISA (BD Biosciences). All samples were diluted 1:16,000 with wash buffer and run in duplicate with average intra- and inter-assay coefficients of variations (CVs) ≤ 10%.

### 2.7. Plasma Endotoxin

Lipopolysaccharide (LPS) was measured in plasma from P21 rat pups by amebocyte lysate method using the Pierce Chromogenic Endotoxin Quant Kit (Thermo). All samples were diluted 1:50 with Endotoxin-Free water and run in duplicate with average CVs ≤ 20% using manufacturer recommendations.

### 2.8. Plasma Inflammatory Cytokines

Detection of inflammatory cytokines: interferon gamma (IFN-γ), interleukin-1β (IL-1β), IL-4, IL-5, IL-6, IL-10, IL-13, chemokine (C-X-C motif) ligand 1 (CXCL1), and tumor necrosis factor-α (TNF-α) was performed in rat plasma samples using MSD V-Plex Proinflammatory Panel 2 Rat kits (Meso Scale Discovery, Gaithersburg, MD, USA). The assay was performed according to manufacturer specifications. All samples were diluted 1:2 and loaded in duplicate with prepared standards onto blocked and washed 96-well plates. Following a series of washes and incubation with antibody detection solution, plates were washed and loaded with read buffer onto a Quickplex SQ 120 Imager [32–38]. Samples reading below the detectable limit of the assay or with coefficient of variation greater than 25% in an individual assay were removed from further analysis [32–38]. The V-Plex pro-inflammatory panel assay is performed with average intra- and inter-assay coefficients of variations (CVs) ≤ 10%. If a replicate was below the curve fit range, it was recorded as 0. Inflammatory marker concentrations are presented as pg/mL.

### 2.9. Short chain fatty acid quantification

Rat stool samples from P21 rat offspring were processed by the Molecular Determinants Core at Johns Hopkins All Children’s Hospital in St. Petersburg, FL. Approximately 100 mg (wet weight) samples of fecal matter were homogenized with 500 ul of 0.5N aqueous HCl in an automated sample mixer (Precellys). After settling of solids, the aqueous phase was transferred to a glass microtube and extracted with 500 µl of methyl tert-butyl ether. A subsample (∼200 µl) was transferred to an autosampler equipped with a glass insert and a PTFE septa cap for analysis via gas chromatography mass spectrometry (GC-MS). Aqueous extracts were re-combined with solids, dried overnight at 60°C, and re-weighed to obtain dry weights for each sample. For GC-MS analysis, 1 µl of each sample was injected onto a 30 meter Agilent FFAP column. Target compounds were quantified alongside isotopically labelled internal standards using selected ion monitoring mode. Limits of quantitation (µg/g dry weight equivalent) were 9 µg/g for acetate, propionate, and butyrate, and 3 µg/g for all other targets.

### 2.10. Western Blotting

Approximately ∼30mg of snap frozen colon tissue was lysed in 1800µL ice cold RIPA buffer with Complete Mini protease inhibitor (Sigma 11836153001) and homogenized using electric homogenizer. Samples were incubated on ice for 4 min and centrifuged at 4°C for 10 min at 14,000 rpm to pellet cellular debris. Supernatant was assayed for protein concentration using the BCA assay (Thermo Scientific 23227). 25µg of denatured protein from each sample was run on SDS-PAGE gel for Western blog using antibodies against Zonula occludens (ZO-1), Occludin (OCLN), Claudin-1 (CLDN1) and the housekeeping gene β-actin (ACTB) for normalization. Proteins were transferred to PVDF membranes and processed for immunoblotting with Li-COR secondary antibodies and imager to quantitate relative protein levels of ZO-1, OCLN, and CLDN1 for each subject by normalizing the signal to ACTB.

### 2.11. Immunofluorescence Staining

All primary and secondary antibodies used in the study are listed in Supplementary Table 2. Colon tissue was collected from rats on P21, and contents were washed out using PBS. Colons were cut longitudinally, laid flat with the luminal side facing up, and gently rolled into a “Swiss Roll” using a toothpick [39]. Rolls were bisected and fixed in 10% Neutral Buffered Formalin for 24 hours prior to tissue processing and embedding in paraffin wax. 5 µm sections were cut using a rotary microtome, mounted onto charged slides, and baked at 55°C for 10 min.

Slides were deparaffinized and rehydrated using a series of xylene and ethanol of descending concentration prior to rinsing with DI water and incubating in fresh 0.2% Sodium Borohydride solution for 5 min at room temperature with mild shaking and washing in TBS. Antigen retrieval was performed by submerging slides in 1x citrate buffer PH 6.0 and steaming for 45 min. After cooling and rinsing, slides were blocked with 5% Donkey serum in TBST for 30 min. Primary antibodies were conjugated to their respective secondary NanoTag antibody as indicated in Supplementary Table 2. The manufacturer recommended concentration for NanoTag secondaries, 20 pmol of secondary Fluotag per ug primary antibody, was used according to the NanoTag Biotechnologies Multiplexing Immunofluorescence (MX-IF) protocol. Each primary was incubated with its respective secondary antibody in separate tubes with 20 µL of PBS per ug of primary antibody for 20 min at room temperature. Slides were then incubated in a humidified chamber at 4°C overnight with the conjugated primary and secondary antibodies diluted in blocking solution. The following day slides were rinsed with DI water and coverslipped with ProLong Gold Antifade mountant with DAPI (Invitrogen P36931). Slides were cured at room temperature for 24 hours and stored at 4°C. Images were captured using a Leica SP8 confocal microscope at 63x magnification, thresholded using the Moments method through ImageJ, then quantified using the Fiji plugin, BIOP-JaCOP, for colocalization between markers [40]. A macro was created to import files with individual channel images into Fiji.

### 2.12. Colon Tissue Histology

Colon tissue was collected from rats on P21, and contents were flushed out with PBS using a syringe and pipette tip. Colons were cut open longitudinally, laid flat with the luminal side facing up, and gently rolled into a “Swiss Roll” using a toothpick [39]. Rolls were bisected and fixed in 10% Neutral Buffered Formalin for 24 hours prior to tissue processing and embedding in paraffin wax. 5 µm sections were cut using a rotary microtome, mounted onto charged slides, and baked at 55°C for 10 min.

Slides were deparaffinized and rehydrated using a series of xylene and ethanol of descending concentration prior to rinsing with DI water. Slides were submerged in Alcian Blue Solution, pH 2.5 (Sigma Aldrich 1.01647.0500) for 5 min and rinsed thoroughly prior to submerging in Nuclear fast red-aluminum sulfate solution 0.1% (Millipore Sigma 60700) for 10 min. Slides were rinsed thoroughly again and dehydrated using a series of ascending ethanols and xylene prior to coverslipping with Permount Mounting Medium (Fischer Scientific SP15-100).

### 2.13. Statistical Analysis

Statistical analysis was completed using GraphPad Prism software (v.10.4.2, GraphPad Software, La Jolla, CA, USA). Data are expressed as averages ± standard error of the mean (SEM). Statistical comparisons were analyzed using factorial or repeated measures 2-way ANOVA with ‘maternal diet’ (Diet) and ‘sex’ (Sex) as independent variables. Post hoc uncorrected Fisher’s least significant difference (LSD) tests were used to conduct pairwise comparisons between diet groups and by sex. The use of any other statistical comparison, such as unpaired t-test or mixed effects models were noted in the results. For all statistical tests, p < 0.05 was considered significant.

## 3. Results

### 3.1. HF offspring have increased body weight gain and plasma leptin

Throughout gestation and lactation, HF-fed dams had significantly increased caloric intake compared to CHOW-fed dams (main effect of diet P < 0.0001) (Fig. 1b). While dams from both diet groups gained weight during gestation, HF-fed dams gained weight more quickly (time*diet P <0.0001) (Fig. 1a). Offspring body weight was similar at birth between maternal diet groups and remained similar until P14 at which point both male and female HF offspring had significantly increased body weight compared to CHOW controls, which persisted until weaning at P21 (main effect of diet: Male P<0.01; Female P<0.05) (Fig. 1c, d). Plasma leptin was significantly increased at P21 in both male and female HF offspring (main effect of diet P < 0.0001), suggestive of increased adipose tissue and leptin resistance (Fig. 1e).

**Figure 1.**
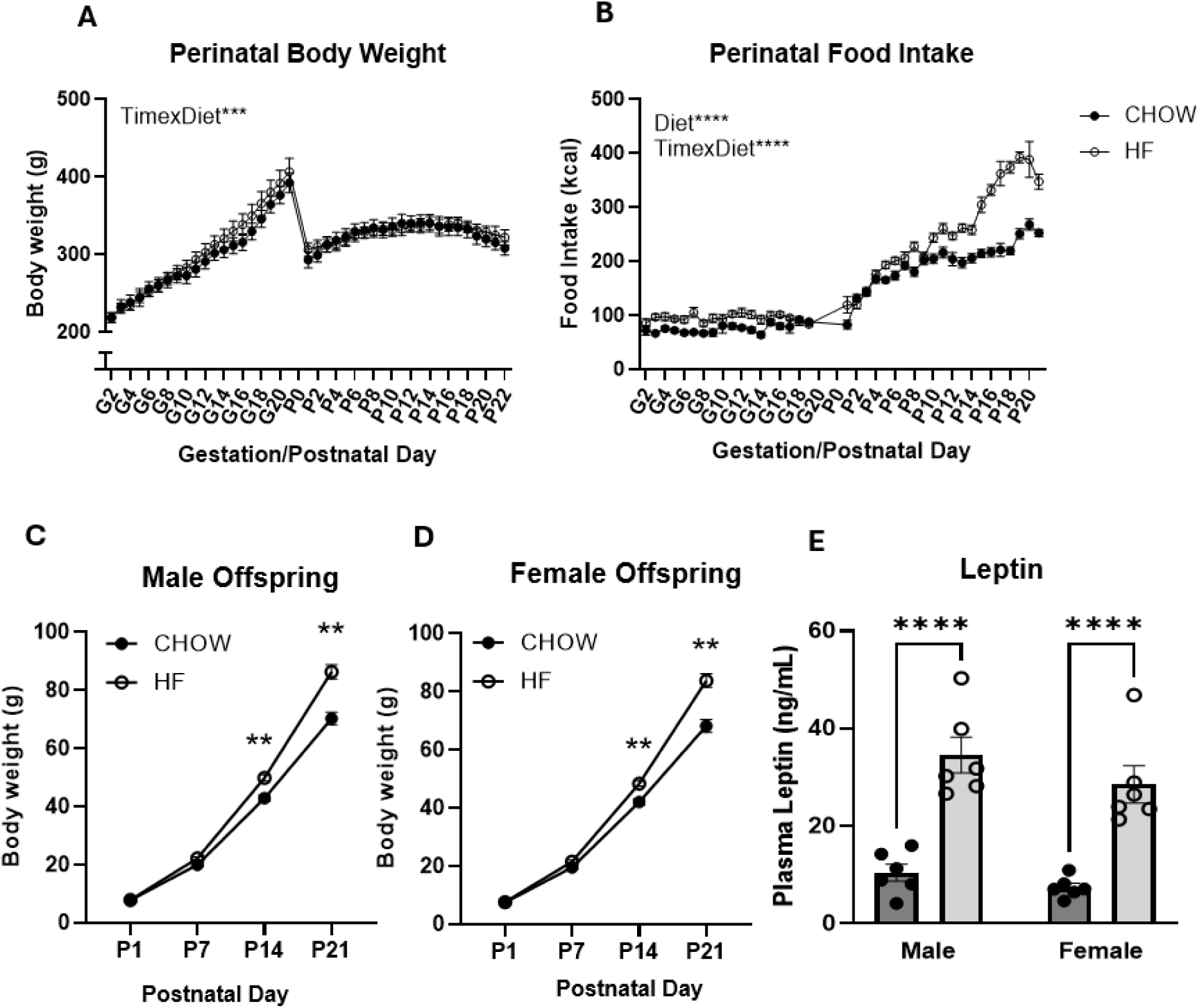
HF offspring have increased bodyweight and plasma leptin at P21. (A) Body weight and (B) Food Intake of pregnant dams during 21 days of gestation. Body weight in postnatal (C) male and (D) female offspring. (E) Male and female plasma leptin concentration on postnatal day 21. All data are presented as mean +/-SEM (n = 8/group). **P < 0.01, ****P < 0.0001 assessed by two-way ANOVA with multiple comparisons. ANOVA, analysis of variance; SEM, standard error of the mean.

### 3.2. HF male offspring have vagal remodeling in adulthood, while females do not

One way in which post-ingestive feedback signaling can be impaired is via remodeling of vagal afferents terminating in the NTS. It has previously been shown that prolonged HFD consumption is sufficient to reduce vagal afferent innervation terminating in the NTS [18]. We tested whether perinatal high fat diet consumption was sufficient to reduce vagal afferent innervation in the NTS. We quantified IB4-positive staining at the level of the medial NTS in adult and P21 male offspring. IB4 selectively binds to unmyelinated C-fibers and can be used as a proxy for density of vagal afferent innervation inputs to the NTS. While there were no significant differences in IB4 staining at P21 (Supplementary Fig. 1a,b), adult male HF offspring had significantly lower IB4 staining (Fig. 2a,c) (P < 0.05), indicative of decreased c-fiber innervation in the NTS in adult male HF offspring by P120. HF female offspring did not show a significant decrease in NTS innervation in adulthood, as seen in male offspring (Fig. 2b,d).

**Figure 2.**
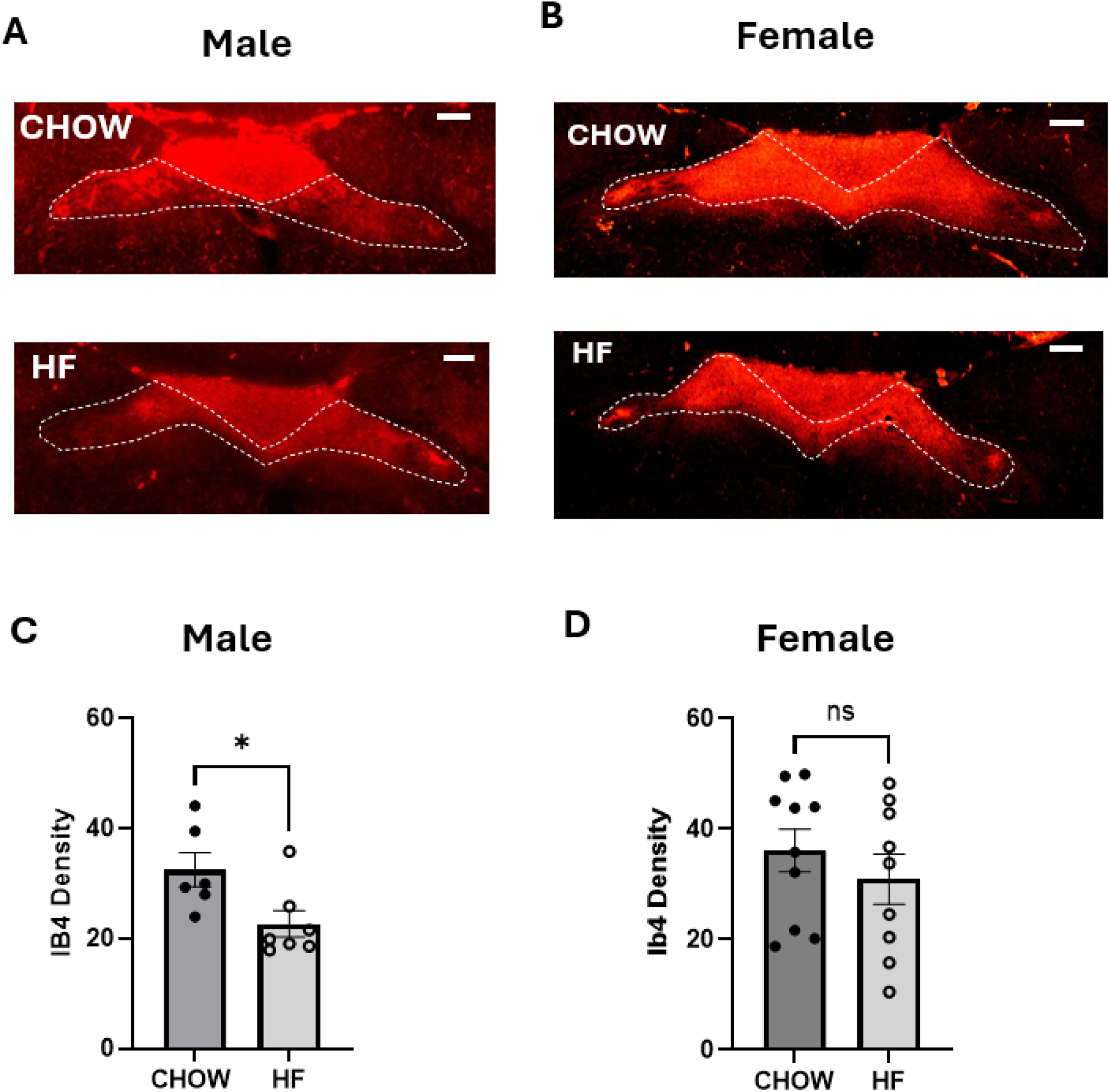
HF male offspring have vagal remodeling lasting into adulthood. Representative NTS sections immunofluorescently labelled for IB4, a marker for c-fiber innervation in adult (A) male and (B) female offspring. (C, D) Average IB4 density measured in the NTS. NTS: nucleus tractus solitarius, IB4: isolectin B4. Data are presented as mean +/-SEM (n = 7-8/group). *P<0.05, assessed by Student’s t-test. Scale bar equivalent to 200uM.

### 3.3. HF male offspring have altered expression of intestinal satiety peptides, while females do not

Another mechanism by which post-ingestive feedback signaling can be impaired is via altered development and function of EECs responsible for secreting postprandial peptide hormones in response to nutrients. To measure this, small intestinal segments were collected at P21 and in adulthood to measure expression of genes encoding feeding related peptides, *Cck*, *Gcg*, and *Pyy,* in male and female offspring.

In male adult offspring, there was an overall effect of maternal high fat diet (P < 0.05) decreasing expression of all three genes, *Cck*, *Gcg*, and *Pyy,* throughout all three segments of the small intestine (Fig. 3a-c), with the HF male offspring having downregulated expression levels for genes encoding satiety hormones when compared to levels in controls. In contrast, there were no significant differences between diet groups in any of these satiety peptides in all three segments of the small intestine in female offspring. This is consistent with our previously published data showing that male and female HF offspring display different metabolic phenotypes in adulthood [8].

**Figure 3.**
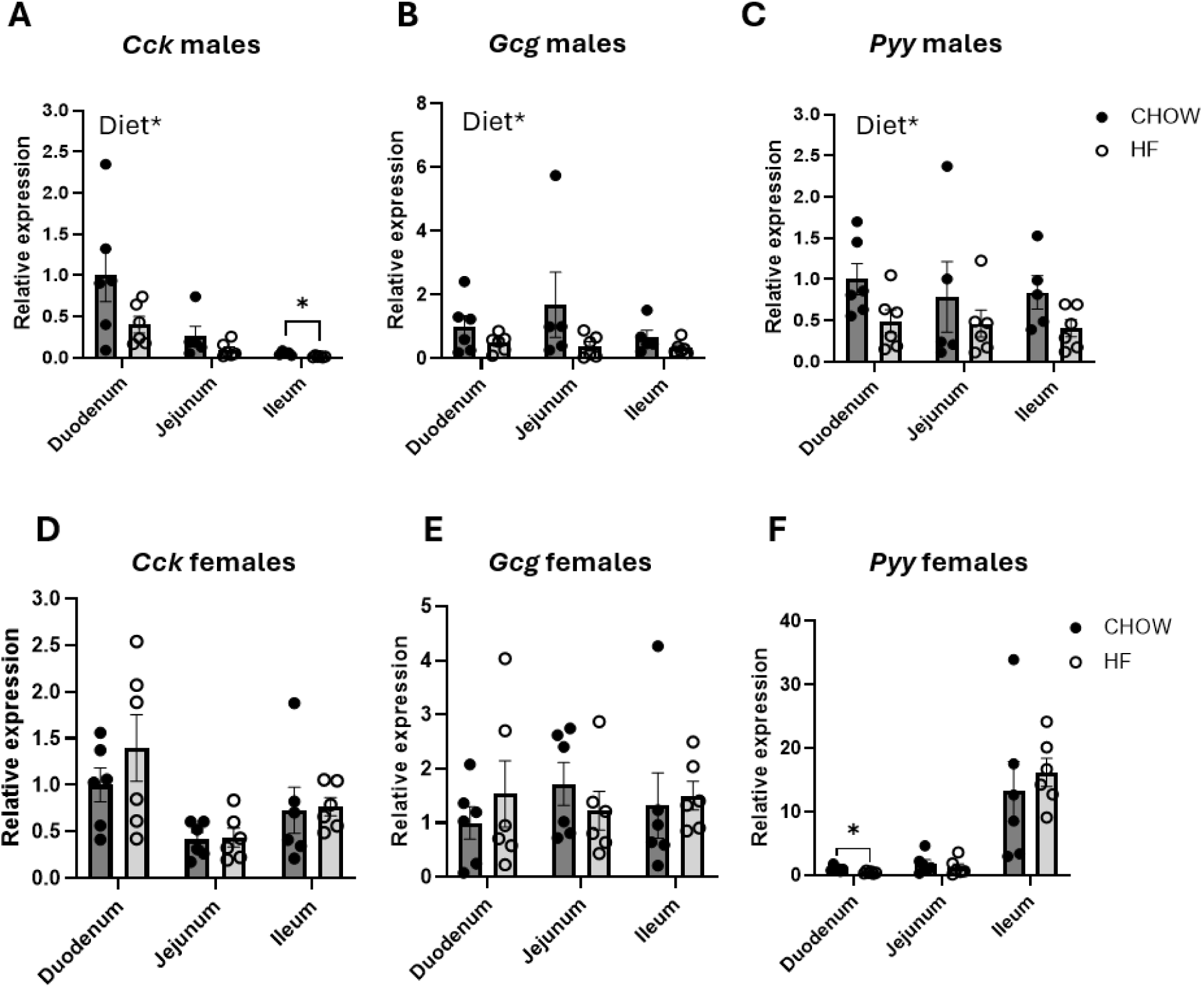
HF male offspring have decreased gene expression of intestinal satiety hormones lasting into adulthood. Male (A-C) and female (D-F) small intestinal tissue collected in adulthood was measured for mRNA expression of (A,D) *Cck*, (B,E) *Gcg*, and (C,F) Pyy relative to *ActB* in the duodenum, jejunum, and ileum. *Cck*: cholecystokinin, *Gcg*: Preproglucagon, *Pyy*: Peptide YY. Data are presented as mean +/-SEM (n = 6/group). *P<0.05, **P < 0.01, **** P < 0.0001 assessed by two-way ANOVA with multiple comparisons. ANOVA, analysis of variance; SEM, standard error of the mean.

At P21, male HF offspring had significantly decreased expression of *Cck* in the duodenum (P < 0.001) similar to the adult offspring (Fig. 4a). *Gcg* and *Pyy* expression were significantly increased in HF male offspring at P21 specifically in the ileum (Fig 4b,c), which differs from the decreased *Gcg* and *Pyy* expression in adulthood (Fig 3b,c). One potential reason for this might be that the pups have access to high fat diet at P21 prior to weaning on to CHOW, and fat in particular is a potent stimulator of GLP-1 and PYY [41]. Similar to the adult data, there were no significant differences in gene expression of *Cck*, *Gcg*, and *Pyy* in P21 females (Fig 4d-f).

**Figure 4.**
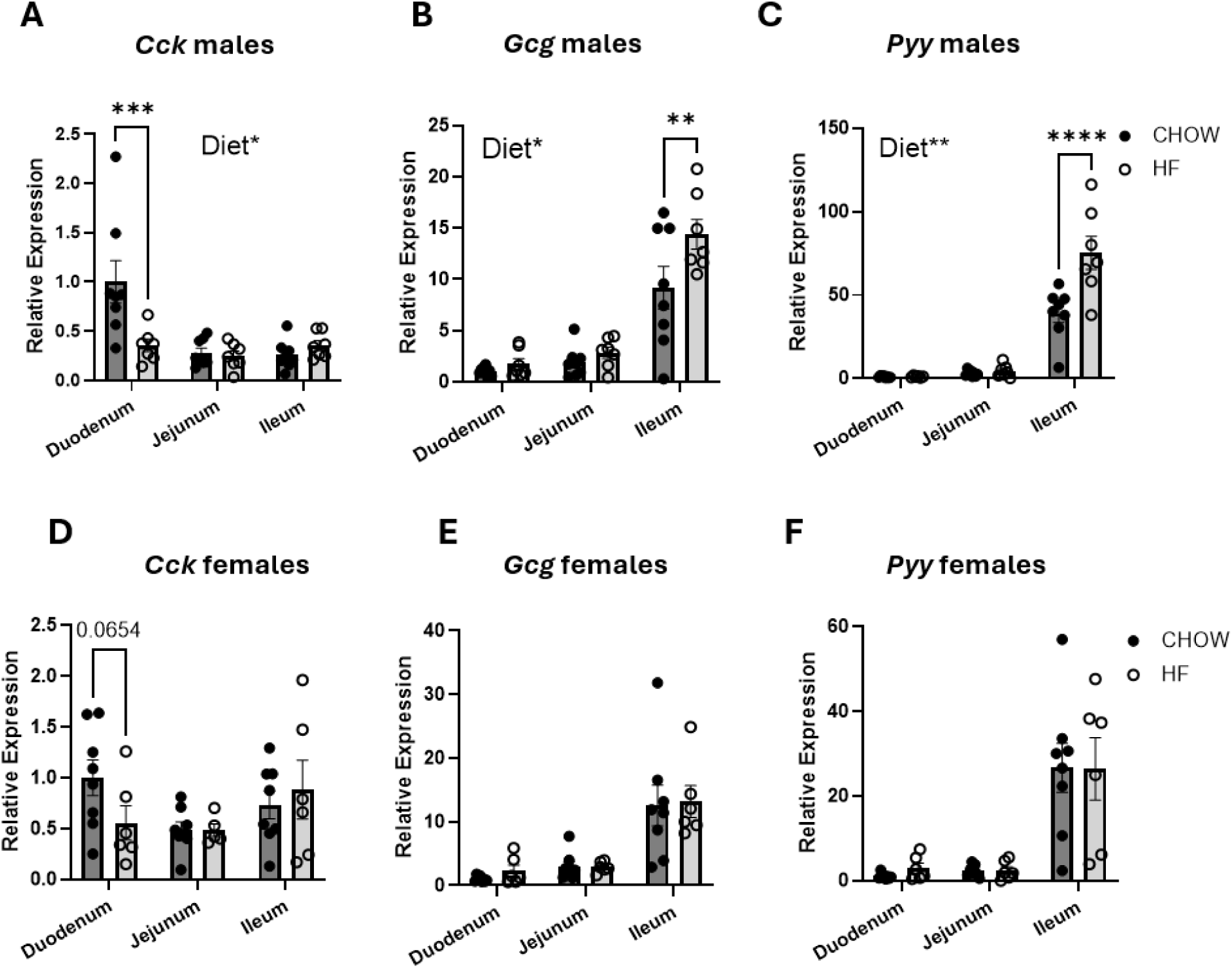
HF offspring have decreased gene expression of intestinal satiety hormones at P21. Male (A-C) and female (D-F) small intestinal tissue collected in adulthood was measured for mRNA expression of (A,D) *Cck*, (B,E) *Gcg*, and (C,F) Pyy relative to *ActB* in the duodenum, jejunum, and ileum. *Cck*: cholecystokinin, *Gcg*: Preproglucagon, *Pyy*: Peptide YY. Data are presented as mean +/-SEM (n = 6/group). *P<0.05, **P < 0.01 assessed by two-way ANOVA with multiple comparisons. ANOVA, analysis of variance; SEM, standard error of the mean.

### 3.4. HF offspring have increased systemic inflammation during development

We hypothesized that inflammation occurring during development may be responsible for alterations in gut-brain axis development resulting in impaired satiety signaling, as evident by altered vagal afferent innervation and expression of intestinal satiety hormones. Obesity has previously been shown to be associated with increased low-grade chronic inflammation [20]. Based on the body weight difference and increased plasma leptin on P21 (Fig. 1c-e), we predicted that other inflammatory markers may also be elevated at this timepoint.

At P21, there was an effect of maternal diet on plasma CRP with HF male and female offspring having increased CRP compared to CHOW (main effect of diet P < 0.01) (Fig. 5a). While there was no significant effect of maternal diet on plasma LPS at the P21 timepoint, there was a sex effect with males having elevated LPS compared to females (sex effect P < 0.01) (Fig. 5b). There was also an effect of maternal diet on IL-1β at P21 with increased levels in HF offspring (main effect of diet P < 0.001) (Fig 5c). There were no significant differences in plasma IL-4, IL-5, or IL-10 although there was a trend towards an increase in IL-5 levels (diet effect P = 0.08). There was a sex effect on plasma IL-6 with female offspring having increased levels compared to males (sex effect P <0.001) (Fig. 5d). Unexpectedly, plasma TNF was significantly decreased in HF offspring compared to CHOW animals (P < 0.05) (Fig. 5f).

**Figure 5.**
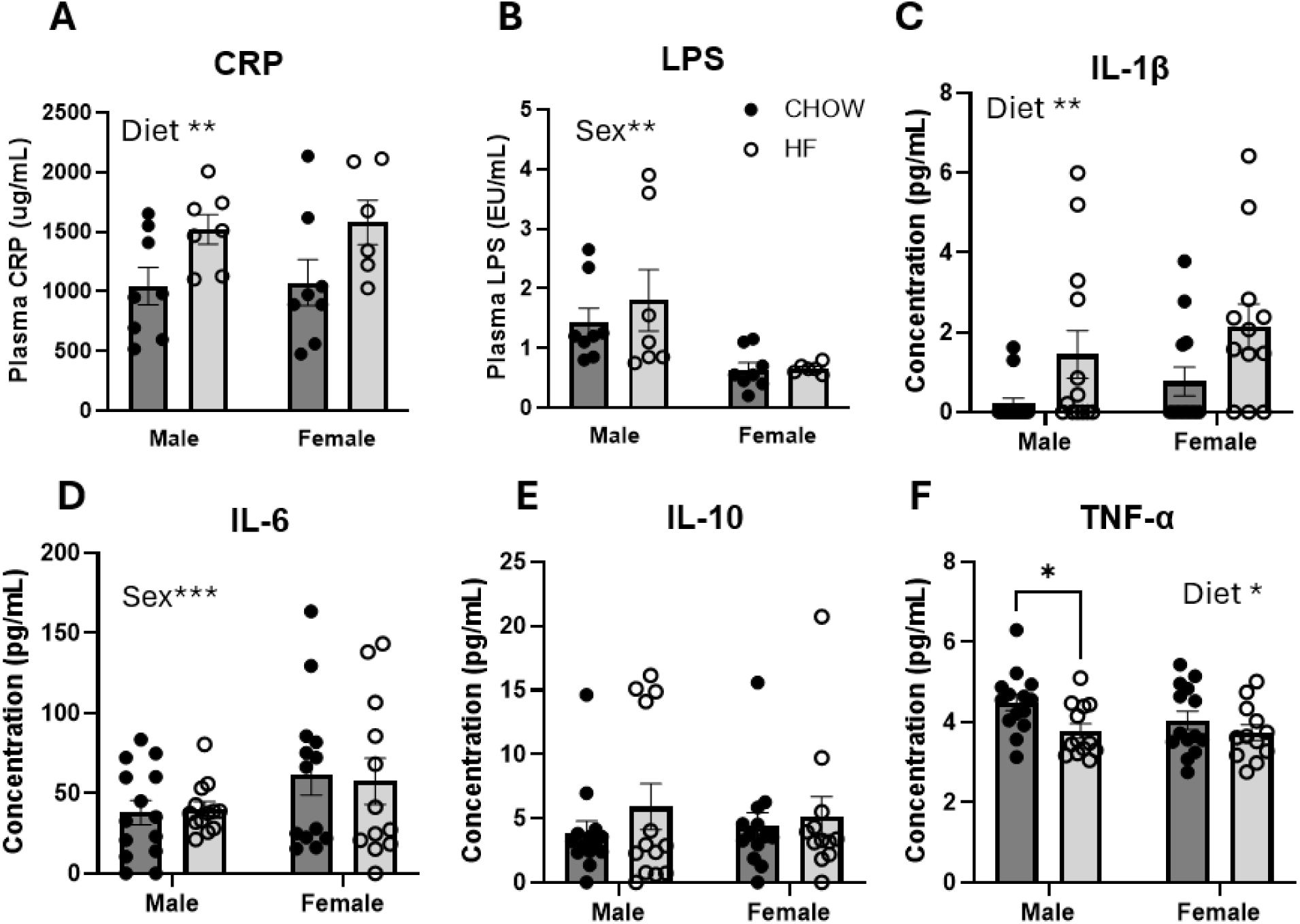
**HF offspring have elevated systemic inflammation during development at the time of weaning**. Concentrations of (A) CRP, (B) LPS, (C) IL-1β, (D) IL-6, (E) IL-10, and (F) TNF-α measured in plasma collected from male and female rat offspring on P21. Chow group is represented by dark circles and the high fat diet group is represented by open circles. Data are presented as mean +/-SEM (n = 6-12/group). *P<0.05, **P < 0.01, **** P < 0.0001 assessed by two-way ANOVA with multiple comparisons. ANOVA, analysis of variance; SEM, standard error of the mean.

### 3.5. HF offspring have altered colonic short chain and long chain fatty acids

We have previously shown using the same perinatal high fat diet paradigm that microbiome composition is significantly altered at P21 [31]. Based on these previous 16s rRNA sequencing data, there were several bacterial taxa that were significantly altered, some of which are producers of SCFAs, namely *Turicibacter* [42] and *Bacteroides* [43], which were depleted in HF offspring. Thus, we predicted that SCFA may be depleted in stool of HF offspring as well. Stool levels of SCFA and LCFA generally correlate with colonic levels [44, 45] and thus were used as an indirect measure of local metabolite production. Stool was collected from P21 pups and processed for analysis by GC-MS. Both male and female HF offspring had significantly depleted levels of the most abundantly produced SCFAs, acetate (P < 0.0001), butyrate (P < 0.05), and propionate (P < 0.01) (Fig. 6a-c). HF male and female offspring also had significantly increased LCFA present in stool compared to CH offspring (main effect of diet P <0.001) (Fig. 6d,e), which may have been a direct result of pups consuming the dam’s high fat diet or increased fat content of the HF dam’s milk composition, which we have previously reported [30].

**Figure 6.**
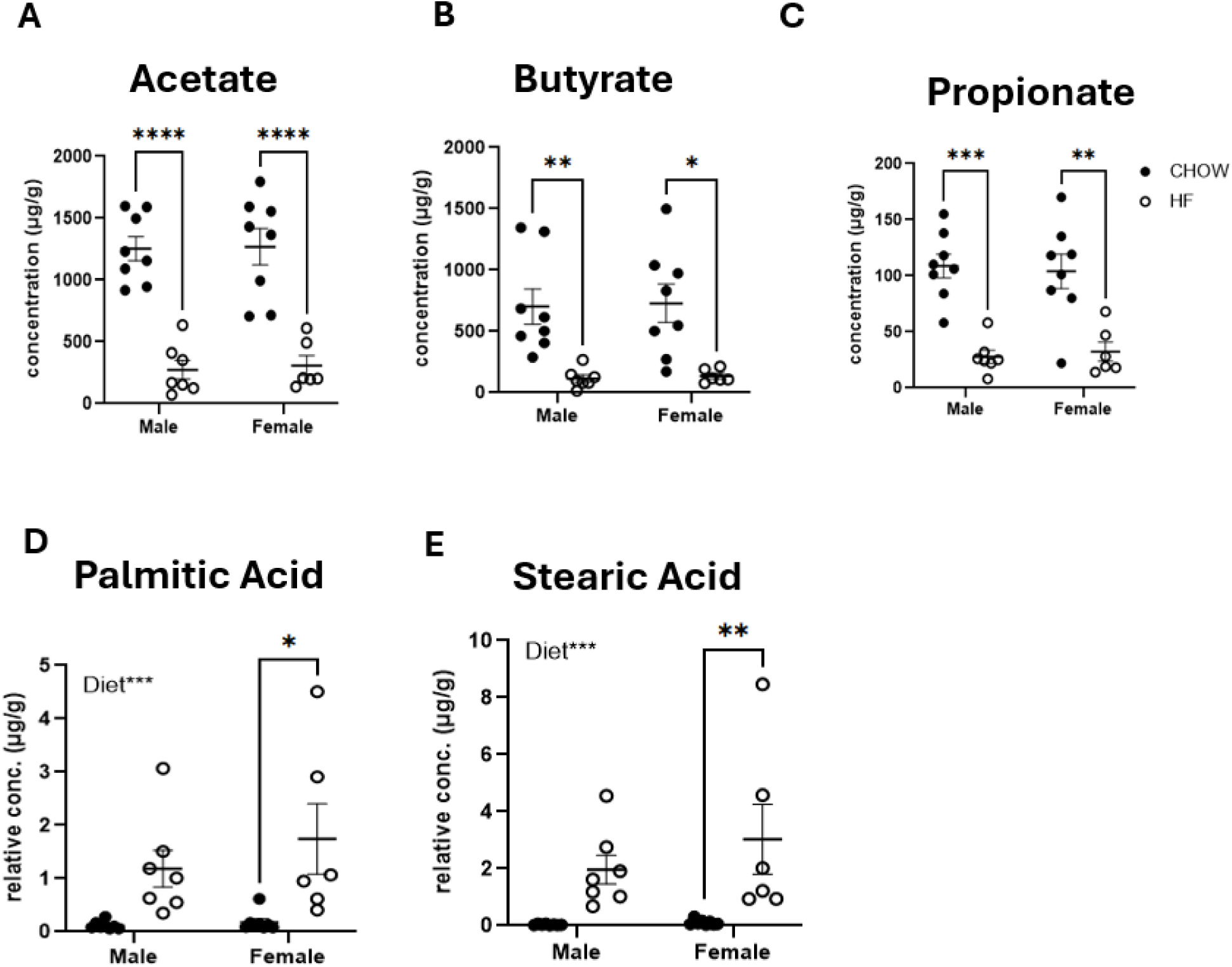
HF offspring have decreased levels of short chain fatty acids present in stool during development and increased levels of long-chain fatty acids at the time of weaning. Concentrations of (A-C) short chain fatty acids, (A) acetate, (B) butyrate, (C) propionate, and (D-E) long chain fatty acids (D) Palmitic Acid, (E) Stearic Acid measured in colon contents collected from male and female rat offspring on P21. Data are presented as mean +/-SEM (n = 6-8/group). *P<0.05, **P < 0.01, **** P < 0.0001 assessed by two-way ANOVA with multiple comparisons. ANOVA, analysis of variance; SEM, standard error of the mean.

### 3.6. HF male offspring have altered organization of tight junction protein complexes

To investigate the role of tight junction protein expression in mediating inflammatory processes, levels of Occludin, Claudin-1, and ZO-1 were quantified by Western Blot in colonic tissue collected on P21. There were no significant differences in the total protein expression of Occludin, Claudin-1, or ZO-1 between maternal diet groups at P21 (Supplementary Fig 2. a-f).

To better determine the functionality of these proteins of interest, we used a multiplex immunofluorescence technique to label each protein with a different probe and measure colocalization of paracellular junctional proteins, Claudin-1 and/or Occludin, with the junctional adaptor protein, ZO-1, which is responsible for crosslinking to the actin cytoskeleton (Fig. 7a-f). There was significantly decreased colocalization between Claudin-1 and ZO-1 (P<0.01) and a trend towards a decrease in colocalization between Occludin and ZO-1 (P = 0.0659) in P21 HF male colon tissue using Mander’s Coefficient, which quantified the area and intensity of overlap between the two proteins of interest (Fig. 7d,e). We specifically chose to use Manders’ Colocalization Coefficient (MCC) as opposed to other common measures of colocalization, like Pearson’s Correlation Coefficient, because we were interested in quantifying the physical interaction and organization between these proteins rather than the ability of one protein to predict or impact the abundance of another.

**Figure 7.**
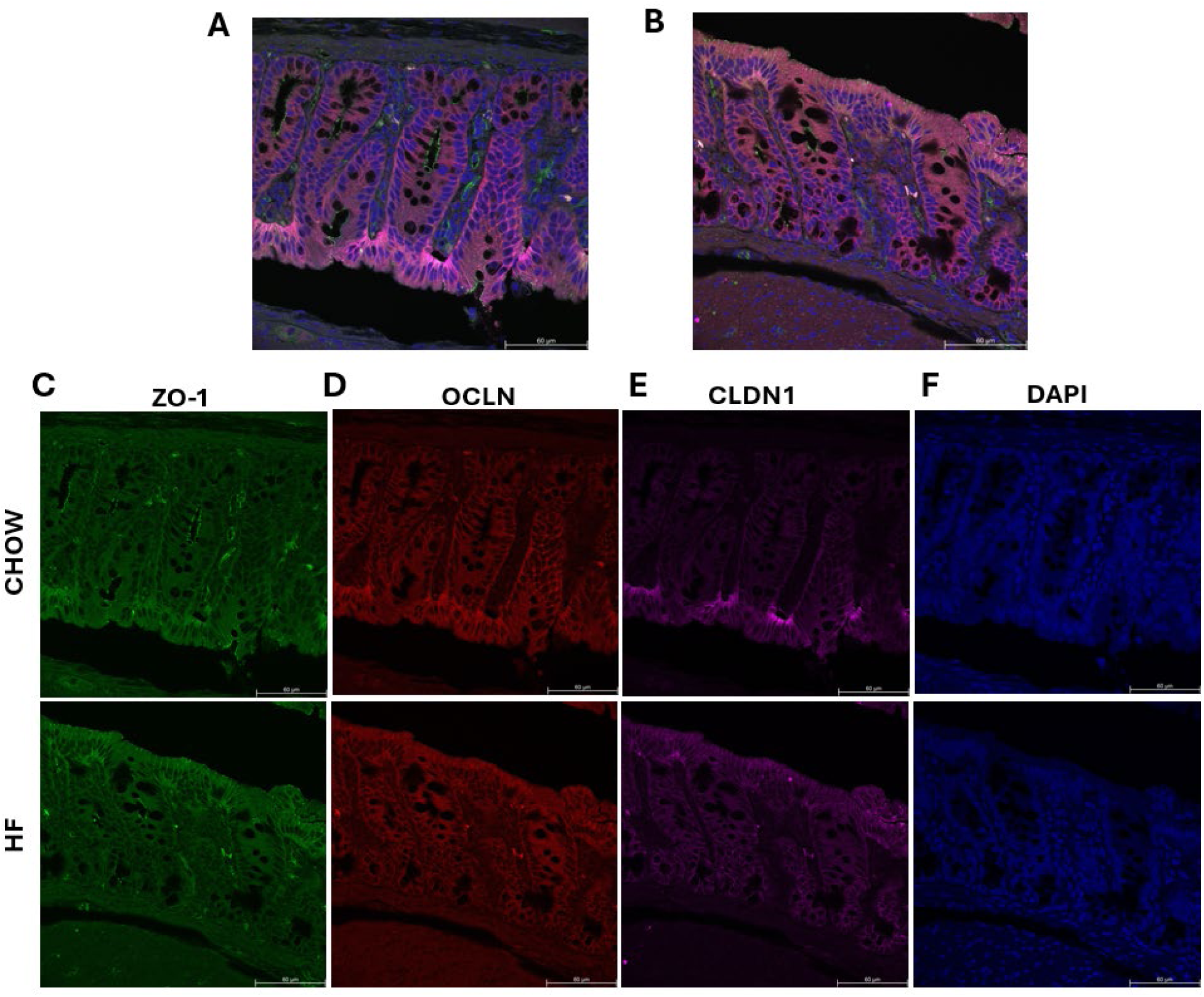
HF animals have decreased colocalization of tight junction proteins measured by immunofluorescent staining. Representative images taken from (A) chow and (B) high-fat offspring colon at P21. Tissue is fluorescently labelled for tight junction proteins (C) ZO-1,(D) OCLN, (E) CLDN1, and counterstained with DAPI. Data are presented as mean +/-SEM (n = 7-8/group). *P<0.05, assessed by two-way ANOVA with multiple comparisons.

In contrast, this decrease in colocalization between tight junction protein markers seen in the HF male colons was not seen in HF females at P21, as there were no significant differences in colocalization between either of the tight junction complexes that were measured (Fig. 7d,e).

### 3.7. HF offspring have altered colonic crypt depth and goblet cell number

H&E staining was performed on P21 colonic tissue to measure histologic changes (Fig. 8a). Both male and female HF offspring had significantly shortened crypt depth at weaning compared to CHOW offspring specifically in the distal colon (P<0.05) (Fig. 8b,c). Additionally, colonic tissue samples were stained using Alcian Blue to look at goblet cell number and with a Nuclear Fast Red counterstain for contrast (Fig. 8d). We used goblet cell number as a proxy for mucus layer thickness, which we had predicted would become thinner in HF offspring at P21. Interestingly, HF offspring had increased goblet cell numbers (Fig. 8e,f), which was contrary to what we had predicted.

**Figure 8.**
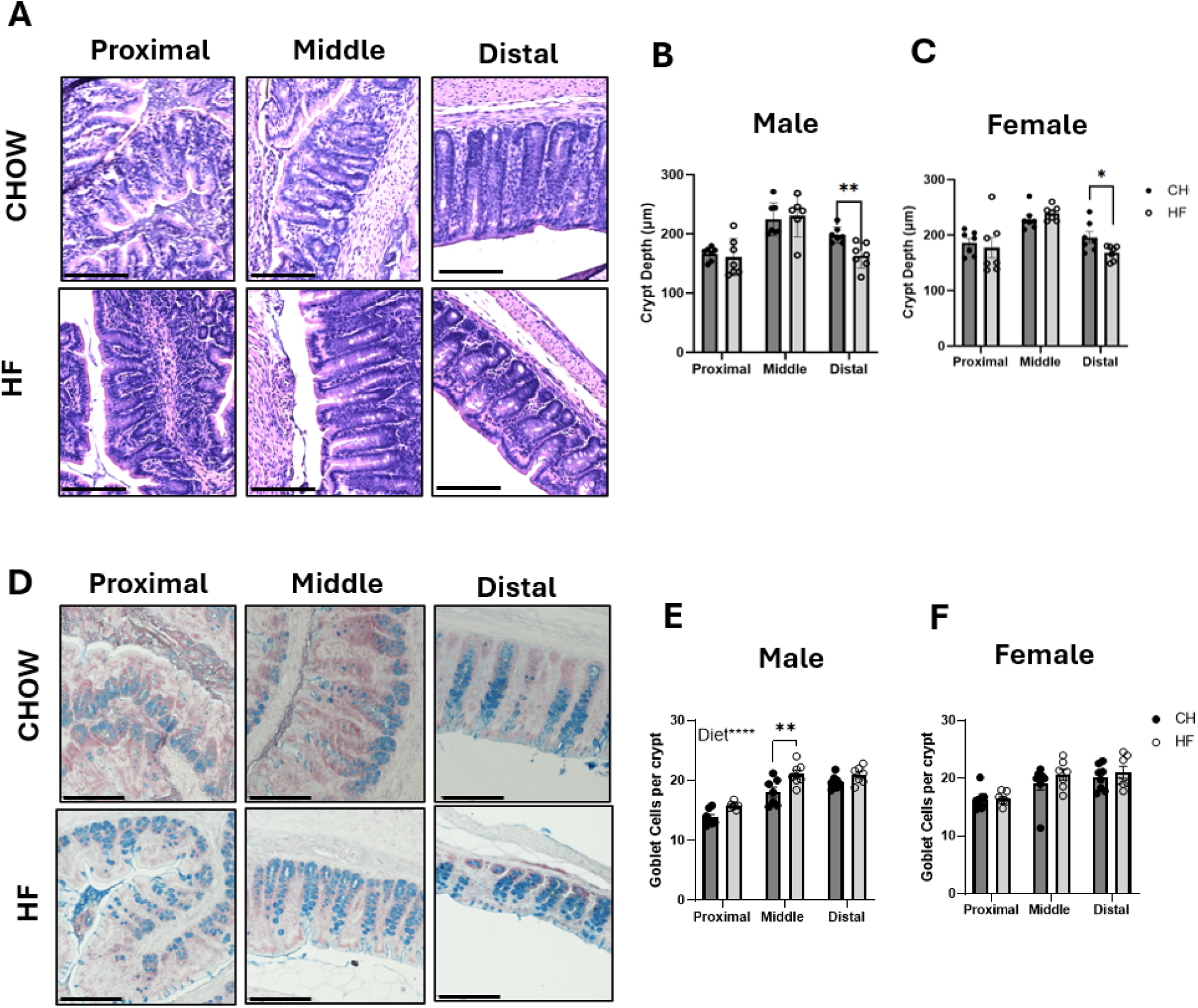
HF offspring have altered crypt depth and goblet cell number during development at the time of weaning. Representative images of H&E (A) and Alcian Blue (D) stained tissue from proximal, middle, and distal colon collected from CHOW and HF offspring on P21. Average crypt depth (B-C) and goblet cell number (E-F) per crypt for male (B,E) and female (C,F) P21 offspring. Data are presented as mean +/-SEM (n = 6-7/group). **P < 0.01, **** P < 0.0001 assessed by two-way ANOVA with multiple comparisons. ANOVA, analysis of variance; SEM, standard error of the mean. Scale bar equivalent to 150uM.

## 4. Discussion and Conclusions

It is well established that prolonged exposure to high-fat diet is sufficient to induce gut dysbiosis and chronic low-grade inflammation, which can result in impaired gut-brain axis signaling [17, 18, 20, 24, 46]. There are several studies that have separately reported maternal HF-diet induced metabolic or immunological impairments in offspring [10, 47, 48]. However, a clear mechanism for increased susceptibility to obesity in HF offspring has not yet been identified. In this study, we report multiple gut-brain axis signaling deficits in male HF offspring, including reduced vagal afferent innervation in the NTS and reduced expression of intestinal satiety hormones. While direct consumption of HF diet has previously been demonstrated to induce vagal remodeling resulting in reduced sensitivity to CCK [18], here we demonstrate the lasting consequences that maternal HF diet consumption can have on offspring, particularly in males.

Similarly, in the small intestine, expression of satiety hormones was also sex-dependent, with adult HF males having reduced expression of *Cck*, *Gcg*, and *Pyy*, while HF females did not. Prior to the beginning of their estrous cycle, P21 females had a trend towards a decrease in duodenal *Cck* expression, while this was no longer apparent in adulthood. The onset of the estrus cycle and hormonal fluctuations in estradiol and/or progesterone may provide metabolic protection against maternal high-fat diet induced obesity. Previous work using ovariectomized and/or postmenopausal females has demonstrated increased susceptibility to HF diet-induced weight gain [49]. Altogether our data suggest that development of the gut-brain axis is impaired in HF male offspring, which may negatively impact post-ingestive feedback mechanisms that drive feeding behavior. These deficits may contribute to the previously reported propensity for obesity in adult male but not female offspring of dams on a HF diet during pregnancy and lactation [8].

During development, we find that HF male offspring have significantly elevated levels of systemic inflammatory markers including plasma CRP, IL-1β, and leptin by P21. We had predicted that plasma LPS may be elevated in the HF offspring due to impaired barrier function as greater intestinal permeability and low-grade chronic inflammation is associated with obesity [20]. Unexpectedly, we found no effect of maternal diet in plasma levels of LPS. There are multiple possible explanations for this finding. Although HF offspring don’t have heightened levels of LPS at baseline, they may instead have a hyper-reactive immune response to similar levels of LPS. In addition, plasma LPS has been shown to have a diurnal rhythm in control animals reaching peak levels at the end of the dark cycle, while LPS levels in HF animals remain high throughout the whole day [24]. Since plasma samples were only collected at one timepoint, it’s possible our LPS data may not be representative of the entire day. To address these possibilities, future experiments may include collecting plasma at multiple timepoints, in addition to isolating immune cells from offspring for LPS stimulation and measurement of inflammatory cytokine secretions. Although there was not a diet effect in plasma LPS, there was a sex effect in plasma LPS with males having increased levels compared to female offspring, which corresponds to our previous finding that HF male offspring are more susceptible to obesity compared to female offspring [8].

Our previous work has identified that HF offspring have altered gut microbiome composition and beta diversity at P21 indicative of gut dysbiosis at this developmental timepoint [31]. From these data, we hypothesized that microbially produced metabolites such as SCFAs may be altered thus impacting inflammatory pathways like NF-KB resulting in increased production of proinflammatory cytokines like IL-1β. As predicted, levels of SCFAs, acetate, propionate, and butyrate, were significantly depleted in stool from HF offspring at P21. These microbial metabolites have been shown to act as immunomodulators, in most cases promoting anti-inflammatory activity, as they are critical for maintaining the integrity of the intestinal immune barrier [50, 51].

Conversely, LCFAs, which were shown to be upregulated in stools of HF offspring, have been shown to promote pro-inflammatory cytokine release when introduced into culture with macrophages [52]. Although we did not measure systemic levels of SCFA and LCFAs in this study, we chose to quantify stool levels as an indirect measure of colonic levels of these metabolites. Further analysis will need to be done to determine whether this translates to changes in systemic levels of these metabolites.

Tight junction (TJ) proteins are critical regulators of intestinal paracellular permeability. Previous literature has shown that prolonged consumption of HFD can promote disruption of intestinal wall integrity by reducing production of TJ proteins resulting in increased translocation of luminal contents, i.e. LPS [20, 46]. We proposed that systemic inflammation in HF offspring at P21 may be due to altered colonic tight junction protein production and/or function. We began by measuring overall total protein content and found no differences between maternal diet groups. To address the functionality of these proteins, we measured the colocalization of TJ proteins using immunofluorescence, and found that male HF offspring had a deficit in colocalization between CLDN1 and the adaptor protein ZO-1, while HF females did not show any differences. ZO-1 is a crucial TJ protein that acts as a scaffolding protein by linking membrane TJ proteins like CLDN-1 and OCLN to the actin cytoskeleton for support. Our data indicate that HF males in particular have a deficit in the organization of these critical TJ proteins, which may be responsible for increased translocation of luminal antigens, potentially resulting in increased release of inflammatory cytokines at this timepoint. This is consistent with the sex effect seen in plasma LPS.

We stained colonic tissue samples to look at morphological changes that may have been induced by perinatal high fat diet and found significant differences in crypt depth and goblet cell number in HF offspring. Previous literature suggests that crypts were evolved in some mammals due to a gradient of microbially derived products such as butyrate, which inhibits stem cell proliferation [53, 54]. The crypt is thought to provide protection for the stem cell niche at the bottom of the crypt from these products. We speculate that decreased colonic butyrate in HF offspring may be linked to their shortened crypt depth. Future work will need to be done to address causation.

Additionally, Alcian Blue staining showed increased numbers of goblet cells present in the mid-colon in HF male offspring, but not females. We intended to use goblet cell number as an indirect measure of mucus production, with the caveat that cell number does address functionality of cells. Previous work shows that long-term high-fat and/or low-fiber diet consumption can lead to increased mucolytic microbial species, which may degrade the mucus barrier [55]. Interestingly, recent work suggests that acute effects of HFD on goblet cell number may differ from long-term chronic effects of high fat diet [56]. One group found in mice that after 10 weeks of HFD consumption, goblet cell density and *Mucin2* expression was significantly increased, while after 16 weeks of HFD consumption, goblet cell number and *Mucin2* expression were significantly decreased, suggestive of a compensatory mechanism to protect from injury to the intestinal barrier [56]. It’s possible that HF male offspring, may have upregulation of goblet cells at P21 as an acute compensatory mechanism similar to these findings that may be followed by eventual disruption to the mucus layer. Further histologic analyses at additional adult timepoints would address whether these consequences persist.

Altogether, these data suggest that perinatal high fat diet induced-gut dysbiosis in offspring results in increased systemic inflammation during a critical period of development, which may contribute to alterations in gut-brain axis signaling, and susceptibility to obesity. Here we have outlined several potential mechanisms that may be responsible for the gut-brain axis deficits seen in adult male HF offspring. Future work aims at finding potential interventions to prevent gut dysbiosis in order to rescue metabolic consequences in offspring.

Additionally, several outcomes had sex-specific effects i.e. NTS innervation, plasma LPS, TJ colocalization, small intestinal expression of satiety hormones, and colonic goblet cell number, while some outcomes occurred in both male and female HF offspring i.e. colonic crypt depth, plasma CRP, and IL-1β. Several outcomes, including gut peptide expression and NTS innervation, were specific to or more altered in male offspring consistent with our previous finding that HF male offspring are more prone to obesity than females in adulthood [8]. This may be due to protective effects estrogen has against dysbiosis and obesity [57–59]. Future work is still needed to address sex differences seen in response to maternal high fat diet exposure.

## Acknowledgments

The authors would like to thank Dr. Cynthia Sears and Dr. Xinqun Wu for their contributions and expertise, the Molecular Determinants Core at Johns Hopkins All Children’s Hospital in St. Petersburg, FL for their technical contributions and expertise in GC-MS measurement of short chain fatty acids. We thank the Phenotyping Core at the Johns Hopkins School of Medicine for their technical services in intestinal tissue processing and sectioning. Supported by NIH DK129193, DK129193-S1, JHU Graduate Student Association Travel Awards to LKM.

