## Supplementary Figures for "Perinatal high-fat diet induces systemic inflammation via altered tight junction complex organization during development and decreases expression of intestinal satiety peptides in male offspring"

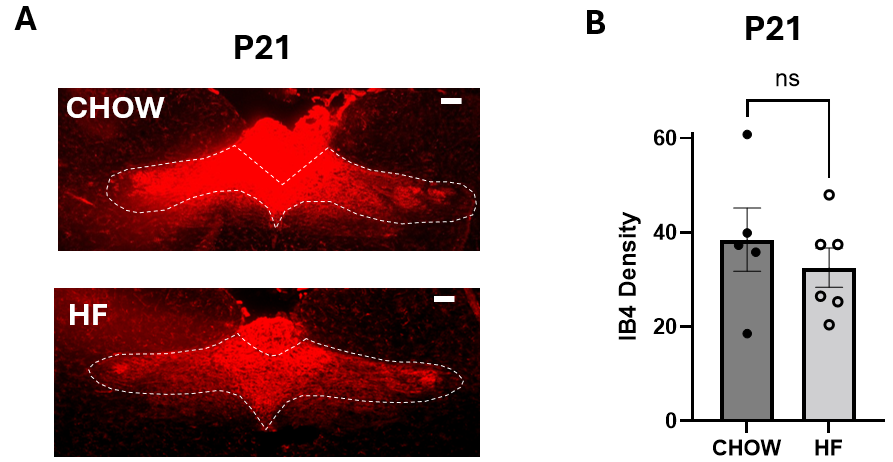

**Supplementary Figure 1.** NTS innervation in P21 male offspring. (A) Representative NTS sections immunofluorescently labelled for IB4, a marker for c-fiber innervation on P21 and (B) average IB4 density measured in the NTS at P21. NTS: nucleus tractus solitarius, IB4: isolectin B4. Data are presented as mean +/- SEM (n = 7-8/group). *P<0.05, assessed by Student’s t-test. Scale bar equivalent to 200uM.

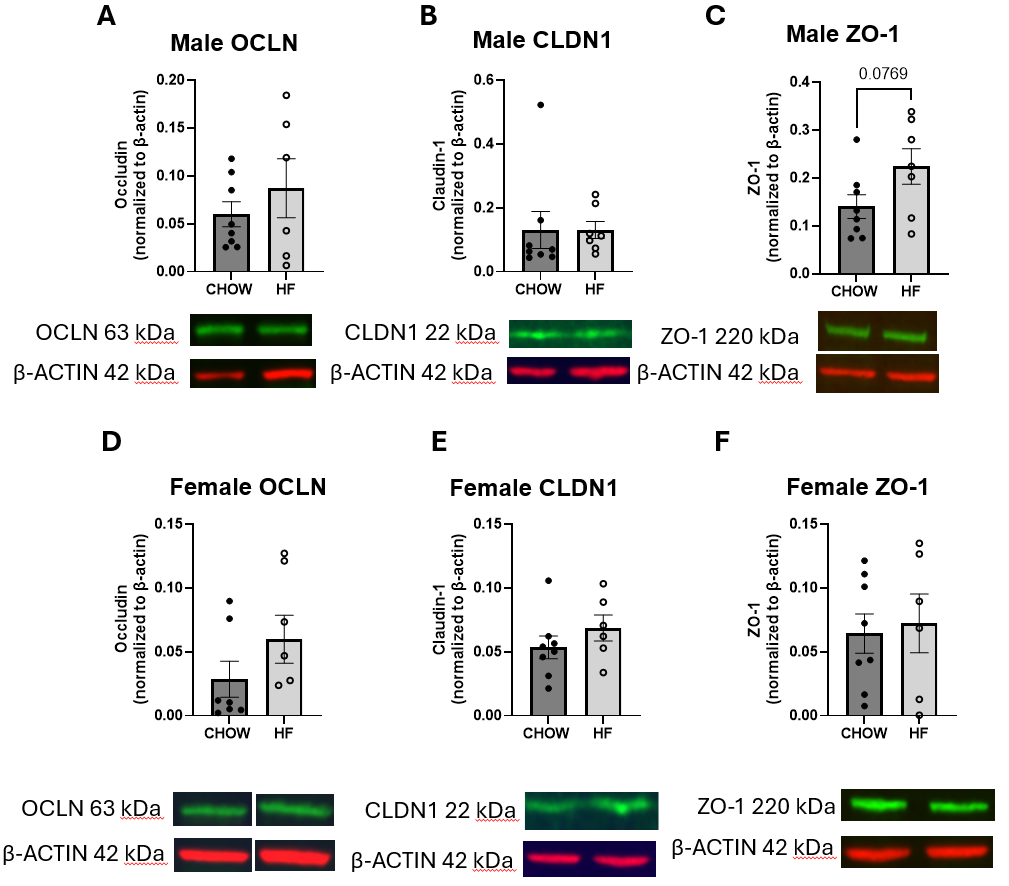

**Supplementary Figure 2.** Western Blot for Tight Junction Proteins in P21 colon tissue. (A-C) Male colon tissue at P21 (D-F) Female colon tissue at P21 (A, D) OCLN/β-ACTIN (B,E) CLDN1/ β-ACTIN (C,F) ZO-1/β-ACTIN

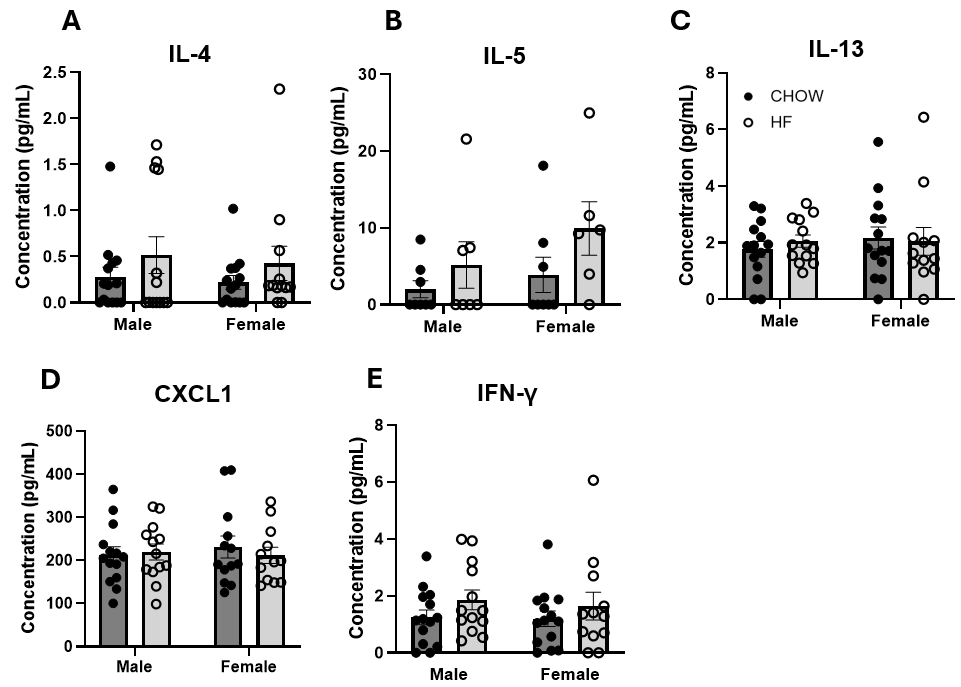

**Supplementary Figure 3. Proinflammatory Cytokines measured in plasma at P21** Plasma concentrations of (A) IL-4, (B) IL-5, (C) IL-13, (D) CXCL1, and (E) IFN-γ measured from male and female rat offspring on P21. Data are presented as mean +/- SEM (n = 6-12/group). Assessed by two-way ANOVA with multiple comparisons. ANOVA, analysis of variance; SEM, standard error of the mean.

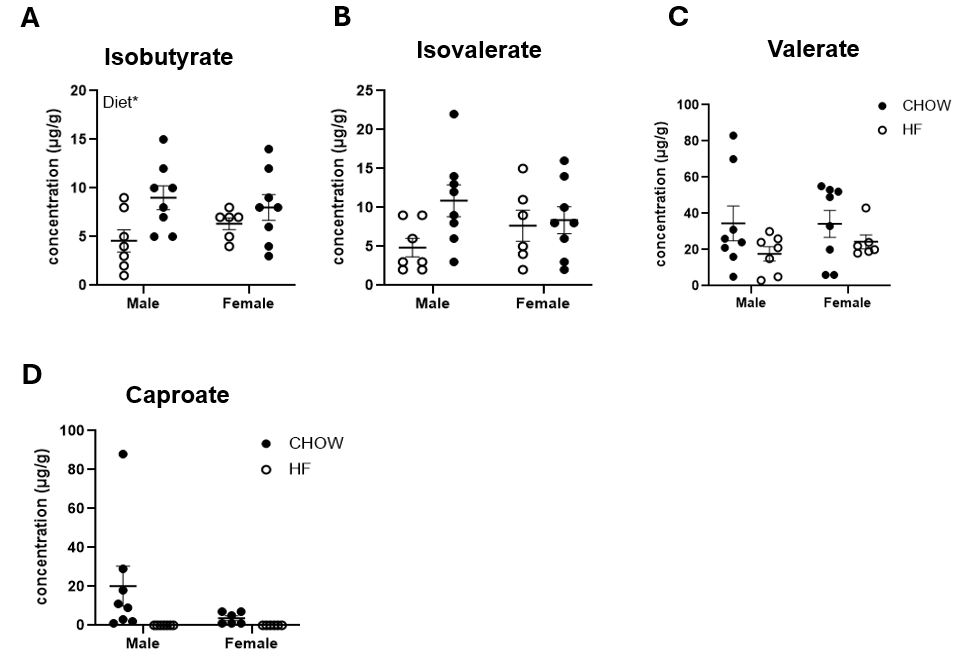

**Supplementary Figure 4. Short chain fatty acid measured in stool at P21.** Stool concentrations of (A) Isobutyrate, (B) Isovalerate, (C) Valerate, and (D) Caproate measured from colon contents taken from male and female rat offspring on P21. Data are presented as mean +/- SEM (n = 7-8/group). *P<0.05 assessed by two-way ANOVA with multiple comparisons. ANOVA, analysis of variance; SEM, standard error of the mean.

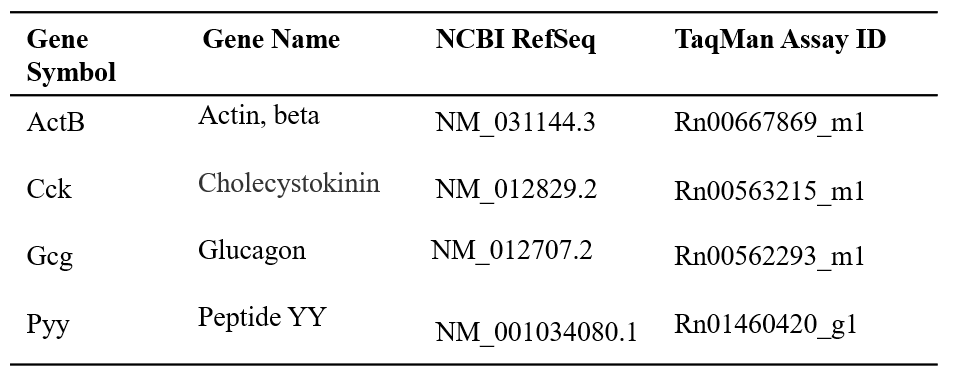

**Supplementary Table 1**. TaqMan assays used for gene expression analysis

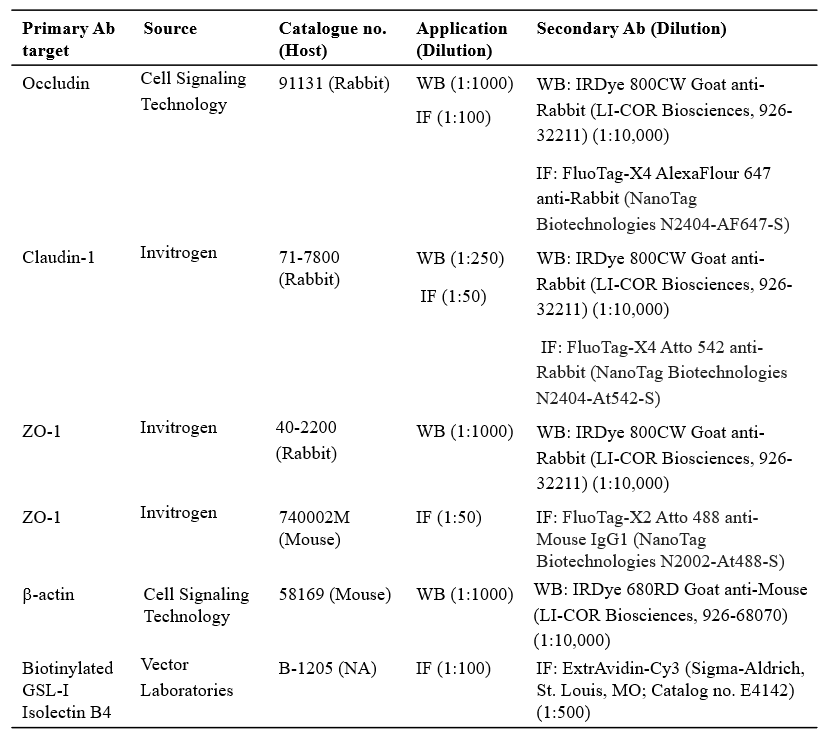

**Supplementary Table 2.** Sources and concentrations of primary and secondary antibodies for immunoblotting or immunofluorescence.
